# Efficient methods for generation and expansion of, and gene delivery to Rhesus Macaque plasma B cells

**DOI:** 10.1101/2021.09.30.462595

**Authors:** Shannon Kreuser, Yuchi Honaker, Rene Yu-Hong Cheng, Noelle Dahl, Rupa Soligalla, Christina Lopez, David J. Rawlings, Richard G. James

## Abstract

Engineered long lived plasma cells have the potential to be a new area of cell therapy. A key step in developing this cell therapy is testing in a model with an intact immune system similar to humans. To that end, we have developed methods to purify, expand, and differentiate non-human primate (NHP; *rhesus macaque*) B cells *ex vivo*. By comparing several media types and conditions, we consistently achieved 10-fold expansion of NHP B cells using a readily available commercial supplement. After only seven days in culture, large percentages of cells in NHP B cell cultures were differentiated. These cells expressed surface markers found in human antibody secreting cells (CD38 and CD138) and secreted immunoglobulin G. We also identified the serotypes (2.5 and D-J) and conditions necessary for efficient transduction of NHP B cells with AAV vectors for the purposes of producing a secreted protein (BAFF). We hope that this work will accelerate proof-of-concept *in vivo* studies using engineered protein-secreting B cells in an NHP model.

## INTRODUCTION

Protein or peptide drugs, including monoclonal antibodies (mAbs), are a growing class of therapeutics that now constitute ^~^10% of the pharmaceutical market^1^. Although protein drugs have great promise due to their ability to specifically target pathways and cell types that have eluded small molecule inhibitors, they have many drawbacks. These drawbacks can include poor solubility, requirement for mammalian or human post-translational modifications, and relatively short half-life’s *in vivo*. Because of these drawbacks, protein drugs are relatively expensive and can be difficult to manufacture. Recently, we developed a cell-based method to deliver protein drugs, which we have successfully tested in immune-deficient mice. To do this, we engineer long-lived antibody-secreting B cells to produce these protein drugs (including specific mAbs)^2^.

We, and others have previously shown that human^1^ and murine^2^ plasma cells can engraft in immune deficient mice. However, although engineered and differentiated murine plasma cells do engraft into immunocompetent murine recipients, they were not detectable more than two weeks post transfer^2^. In contrast, engrafted engineered antigen-specific memory B cells exhibit remarkable stability in the immunocompetent murine recipients; these cells were detectable following re-immunization for up to nine months^3,4^. There are many possible reasons for the limited longevity of engineered plasma cells in the murine system including, among other possibilities, the source of the B cells, species differences and the method of differentiation. Because delivery of engineered plasma cells to immunocompetent recipients is a key proof-of-concept prior to advancing a plasma cell-based therapy into a clinical setting, we sought to develop reagents for studies in the non-human primate, *rhesus macaque*.

Here, we identified a series of reagents for isolation, transduction, expansion, differentiation and qualifying engineered B cells taken from peripheral *rhesus macaque* blood mononuclear cells. We found that an optimal growth media for expansion and differentiation of *rhesus macaque* B cells is a commercial reagent developed for use in human B cells. Contrary to our expectations, *rhesus macaque* B cells differentiated into antibody secreting plasma cells at a faster rate in culture than human B cells. Despite this faster kinetic, we predicted that the degree of B cell expansion in this system will facilitate multiple transfers of plasma cells at the same body weight proportions used previously in murine studies of human and murine plasma cell longevity.

## RESULTS

### Culture with a commercial culture media promoted efficient expansion and differentiation of NHP B cells *ex vivo*

Human cytokines and human-directed antibodies may not efficiently cross-react with NHP proteins. To begin study of NHP B cells, we empirically tested these reagents using rhesus macaque B cells. The first step in ex vivo B cell culture and expansion is isolation of cells from peripheral blood mononuclear cells (Figure 1A). We investigated commercial isolation kits that use antibodies that bind sequences conserved between rhesus macaque and homo sapiens (Figure S1) and identified an NHP CD20 positive selection kit that enabled us to purify B cells at >75% purity (Figure 1B).

**Figure 1.**
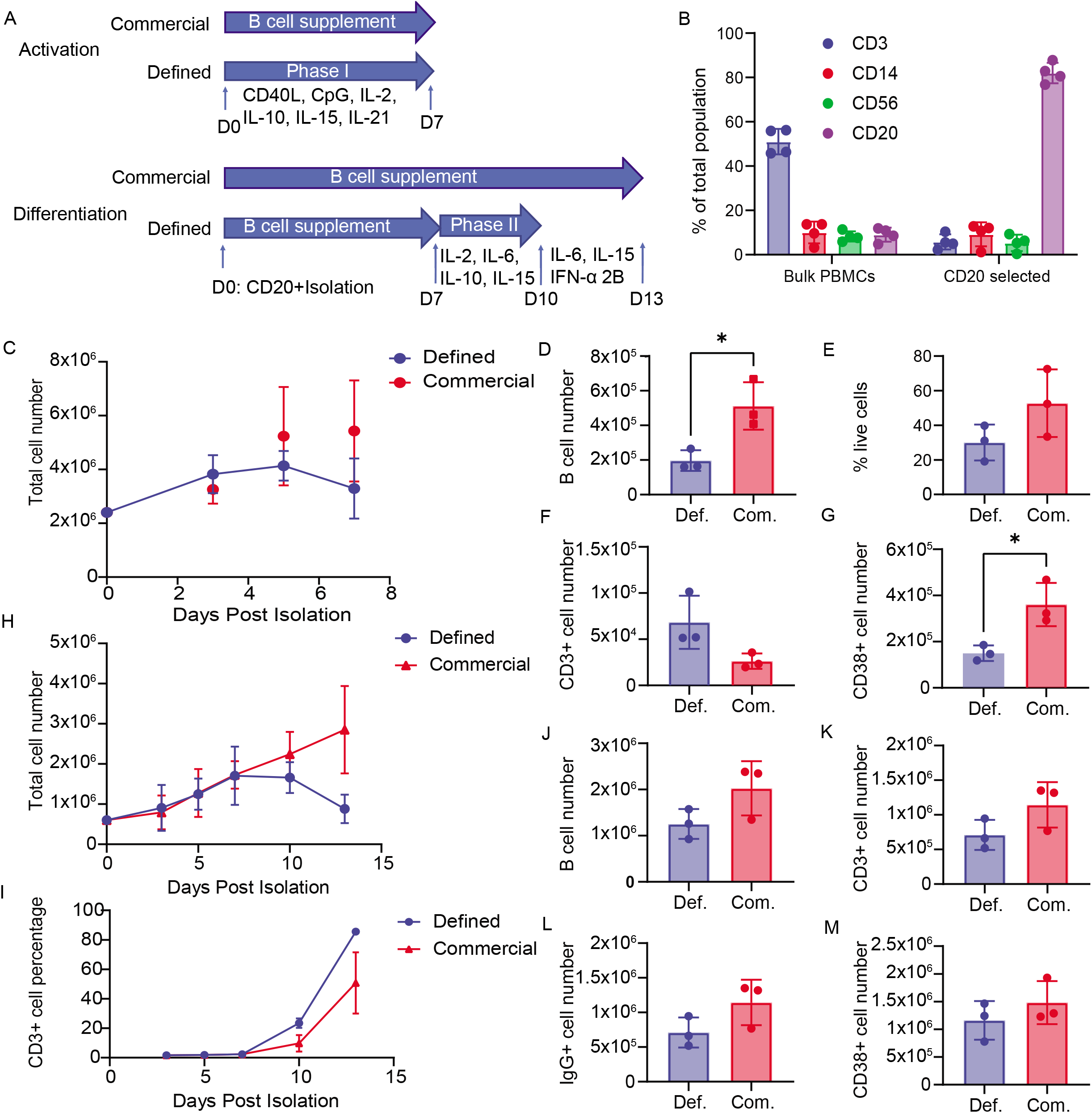
(A) Diagram of experiments comparing commercial and defined culture protocols for either activation (1) or differentiation (2). (B) Following isolation of CD20^+^ Rhesus B cells from PBMCs, we assessed cell purity via flow cytometry using the indicated antibodies (4 donors, n=4). (C-G) CD20^+^ NHP B cells were cultured for 7 days with defined or commercial expansion medium (1.5-1×10^6^ cells/mL)(3 donors, n=3). (C) Total cells were counted on days 3, 5, and 7 (3 donors, n=3). (D-G) On day 7, we used flow cytometry to quantify the number of B cells (D; CD3^-^CD14^-^), viability (E; AF350^--^), T cells (F; CD3^+^) and plasmablasts (G; CD38^+^). (H-M) To assess differentiation in the two protocols, CD20^+^ NHP B cells were cultured (1.5-1×10^6^ cells/mL) in the commercial medium for 7 days, and then cultured for an additional 6 days using either the commercial medium or the defined protocol (bottom panel, A) (3 donors, n=3). (H-M) Total cells were counted (H), and the cells were analyzed by flow cytometry at the indicated time points. We plotted the percentage of contaminating T cells cells (I; CD3^+^), and number of B cells (J; CD3^-^CD14^-^), T cells (K; CD3^+^), IgG^+^ B cells (L) and plasmablasts (M; CD38^+^) on day 10. To assess significance, we used a unpaired t-test (* p < 0.05).

We previously described a method for expanding and differentiating human B cells into plasma cells, which consisted of culture with defined cytokines that activate naive/mature B cells (Figure 1A, Phase I cytokines), followed by culture with cytokines that promote differentiation (Figure 1A, Phase II/III cytokines) ^1^. Here, we compared the activation of isolated NHP B cells *ex vivo* using a modified version of this defined cocktail with a commercially available human B cell expansion cocktail. Following isolation, rhesus B cells were grown in either the defined or commercial medium. During the first seven days of culture, we observed significantly more expansion of NHP B cells grown in the commercial culture, relative to cells grown in the defined media (2.5-fold expansion, Figure 1C-D; gating Figure S2-3). To determine the mechanistic basis for these differences in cell expansion, we quantified cell proliferation and viability. While the percentage of cells proliferating at day 3 in culture (Propidium Iodide; Figure S4) was unchanged between conditions, the viability of B cells grown in the commercial cocktail was increased (Figure 1E). Additionally, we found that culture in the defined medium increased the proportion of CD3+ T cells by day seven (Figure 1F), presumably because of the high concentration of IL2 used in the defined medium. Finally, we found that the commercial media significantly increased the number of cells expressing CD38 (Figure 1G), a marker of differentiation, in CD3-CD14-B cells. These data indicated that the commercial cocktail efficiently activated NHP B cells *ex vivo*.

We next looked at differentiation in the NHP B cells. All cells were cultured in the commercial cocktail for seven days, and then for six additional days using the defined or the commercial cocktails (Figure 1A, bottom panel). In both conditions, total cell numbers decreased (Figure 1H), and the percentage of contaminating CD3+ cells increased (Figure 1I) between days 10 and 13. After 10 days in culture, the commercial cocktail exhibited fewer T cells (Figure 1K), more B cells (Figure 1J) and more IgG+ B cells (Figure 1L) without altering the number of CD38+ B cells (Figure 1M). Collectively, these data demonstrated that NHP B cells cultured in the commercial cocktail for 10 days exhibited optimal expansion and differentiation into IgG+ B cells.

### Low seeding density improves NHP B cell expansion and differentiation *ex vivo*

Cell to cell contact and seeding density play a large role in B cell expansion and differentiation^5,6^. To test the effect of seeding density on NHP B cell cultures, we cultured cells for seven days in the commercial expansion cocktail at high (1.5 million cells mL^-1^) and low (100 thousand cells mL^-1^) densities. We found that NHP B cells grown at low density *ex vivo* exhibited significant increases in total number relative to those grown at high density (Figure 2A-B; ^~^10.5-fold compared to 1.5 fold). We found that B cells grown at low density also exhibited increased viability (Figure 2C) and increased purity as defined by the CD14^-^CD3^-^percentages (Figure 2D; gating Figure S5). Upon comparing the base media formulations, we found that culture of NHP B cells in commercial media plus FBS led to increases in the numbers of B cells relative to culture in ICXF with proprietary supplement (Figure S6).

**Figure 2.**
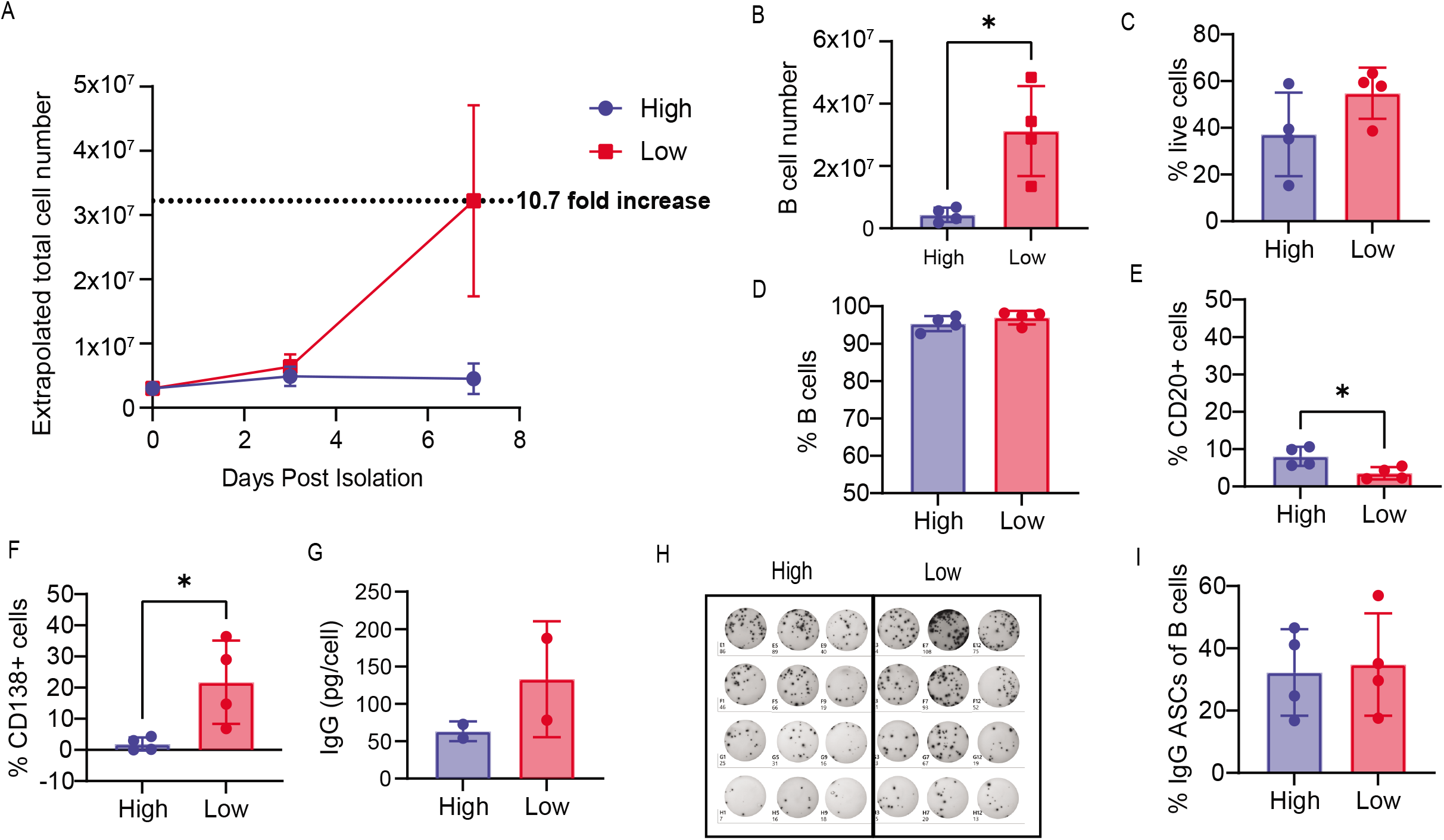
(A-I) To assess the impact of cell density, CD20^+^ NHP B cells were cultured in the commercial B cell medium for 7 days(Plated and maintained 1.5×10^6^ cells/mL versus plated at 2.5×10^5^ cells/mL and maintained at 1×10^5^ cells/mL; 3 donors, n=3). (A) A graph showing the extrapolated total cell numbers for both conditions. At day 7 following isolation, the cells were analyzed using flow cytometry. We plotted the extrapolated number of B cells (B; CD3^-^CD14^-^), and the percent viability (C; AF350^-^), B cells (D; CD3^-^CD14^-^), naive B cells (E; CD20^+^) and plasma cells (F; CD138^+^). (G-I) At day 7, supernatants and cells from the above experiment were assessed for IgG secretion via ELISA (G) and ELISpot (H; image, I; quantification).

Contrary to our expectations, we observed very low percentages of CD20^+^ (Figure 2E) and high percentages of CD138^+^ in NHP B cell cultures (Figure 2F) following only seven days in culture. These phenotyping data indicated that NHP B cells exhibited more rapid differentiation than we previously observed in human B cell cultures^1^. Additionally, we found that culture of NHP B cells at low density resulted in decreased numbers of CD20+ (Figure 2E) and increased numbers of CD138+ (Figure 2F) B cells, implying that the cells cultured at low density differentiated more efficiently. To confirm that the NHP B cells were differentiated into plasma cells, we quantified secreted IgG (Figure 1G) and detected antibody secretion, but at higher levels in low density cultures. Finally, upon plating B cells from each condition, we determined that the percentage of antibody secreting cells detected in ELISPOT was similar (Figure 2H-I). In conclusion, we found that NHP B cells cultured at low density expand and differentiate more efficiently than those cultured at high density.

### Efficient transduction of NHP B cells with the AAV serotypes 2.5 and D-J

To assess the potential for transduction of NHP B cells with recombinant AAV vectors, we isolated B cells, cultured briefly and exposed the cells to a panel of self-complementary AAV vectors. We assessed a series of AAV with alternative serotypes, each carrying a GFP expression cassette to permit efficient tracking of transduced cell populations (Figure 3A). Following exposure to AAV, B cells were maintained in culture for nine more days prior to assessment of GFP positivity in CD3-CD14-cells. In a volume matched experiment (20% AAV by volume), we found that AAV 2.5 and AAV D-J serotypes exhibited the highest percentage transduction (^~^25%; Figure S7A), although the titers varied widely (Figure S7B). To rule out the impact of titer, we directly compared transduction with the 2.5 and D-J serotypes using the same titers. We found that 2.5 and D-J exhibited similar transduction efficiencies, and contrary to our expectations were maintained at high percentages at late time points (Day 10; Figure 3B). Next, to determine whether AAV D-J is useful for delivery of a candidate, physiologically relevant, payload in NHP B cells, we transduced NHP B cells with single-stranded AAV expressing the cytokine BAFF cis linked to GFP (Figure 3C). While the transduction rates using the BAFF AAV vector were slightly lower than the self-complementary GFP vector (Figure 3D), we detected high levels of BAFF secretion (Figure 3E) in NHP B cells. We also tested whether transduction was different in B cells cultured at different densities. We found that NHP B cultured at low density were transduced at higher rates than those cultured at high densities (Figure 3F), and produced more BAFF in culture (Figure 3E). In conclusion, because AAV DJ can reproducibly be produced at higher titers (example Figure S5B), we predict that the DJ serotype will be more useful than 2.5 for transduction of NHP B cells.

**Figure 3.**
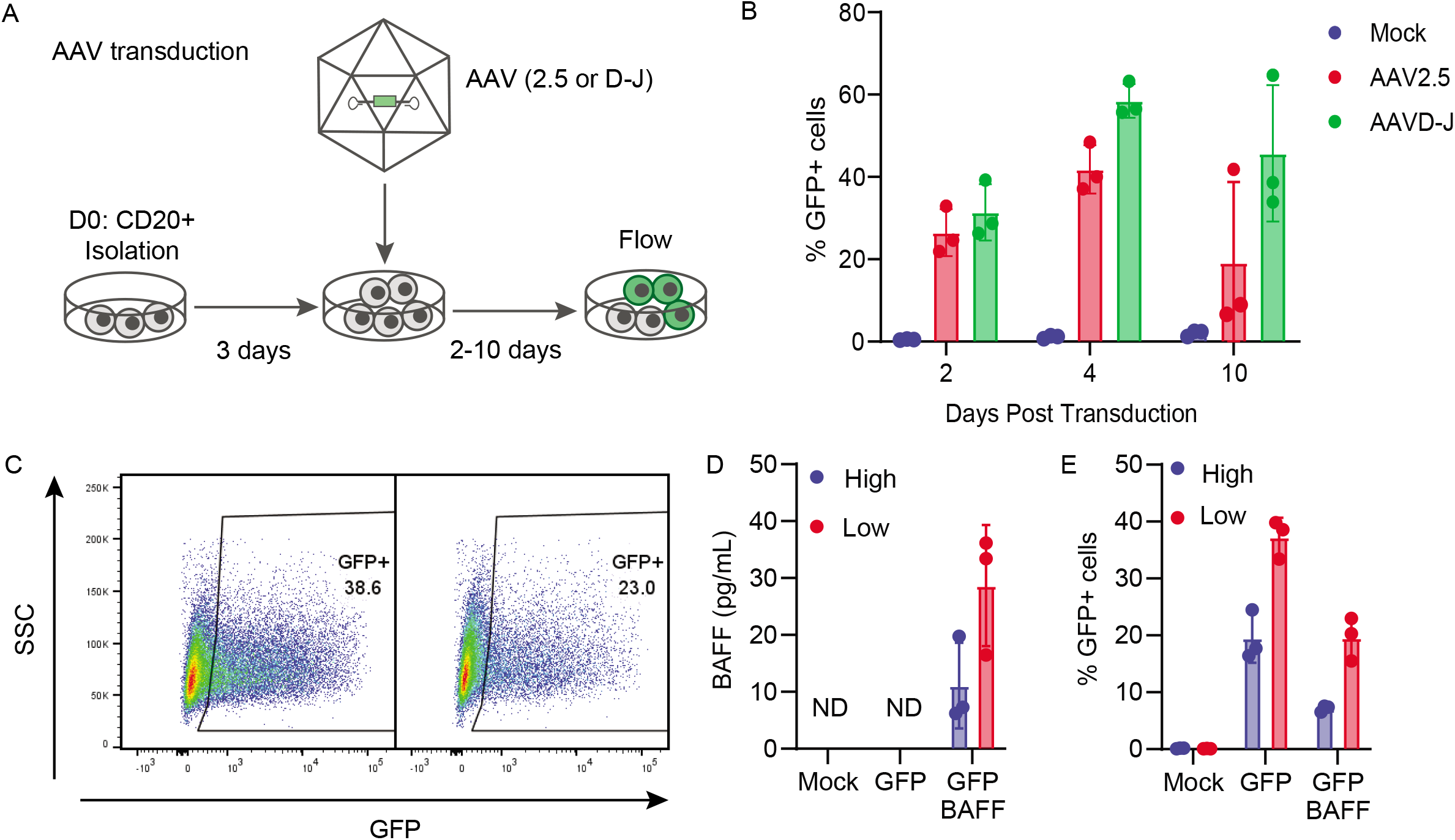
(A) Diagram of the AAV and transduction and culturing protocol. (B) NHP CD20+ B cells were cultured for 3 days before transduction with the indicated AAV pseudotypes expressing GFP (MOI = 4×104). Transduction efficiency was quantified by flow cytometry at the indicated time points. (C-F) NHP CD20^+^ B cells grown at high (1.5×106 cells/mL) or low density (2.5×105 cells/m) were transduced with either AAVD-J GFP or AAVD-J GFP-BAFF (3 donors, n=3). GFP percentage (C, D) and BAFF secretion (E) was quantified four days post-transduction.

### Assessing Lentiviral transduction of NHP B cells for possible use for gene delivery

B cells from multiple species, as well as *rhesus macaque* cells from several hematopoietic lineages, have been historically difficult to transduce with gamma-retroviral or lentiviral vectors^7^. Using the XHIV/SHIV packaging system^8^, we made both VSVg and Cocal pseudotyped lentiviral vectors with a GFP payload. After 2 days in culture, each vector was added to NHP B cells (MOI 40). Following vector delivery, B cells were cultured for 11 days in the commercial media, and the transduction was quantified using flow cytometry. At no point did we observe greater than 2% of the population expressing GFP with either vector (Figure 4B). We also tested lentivirus pseudotyped with a modified version of the Nipah F and G proteins that specifically binds to the CD20^9^ receptor, which is highly expressed on NHP B cells. When using this modified Nipah pseudotype, we found that the percentage GFP positive cells was also less than 3% (Figure 4C), although the mean fluorescent intensity of the GFP positive population was substantially greater than that seen with the VSVg and Cocal pseudotypes(Figure 4C). These results demonstrate that lentiviral transduction of NHP B cells remains a hurdle. However, our findings suggest that using a receptor-targeted Nipah pseudotype that is more specific to NHP CD20 or to a different receptor highly expressed on NHP B cells ultimately could facilitate efficient transduction of NHP B cells.

**Figure 4.**
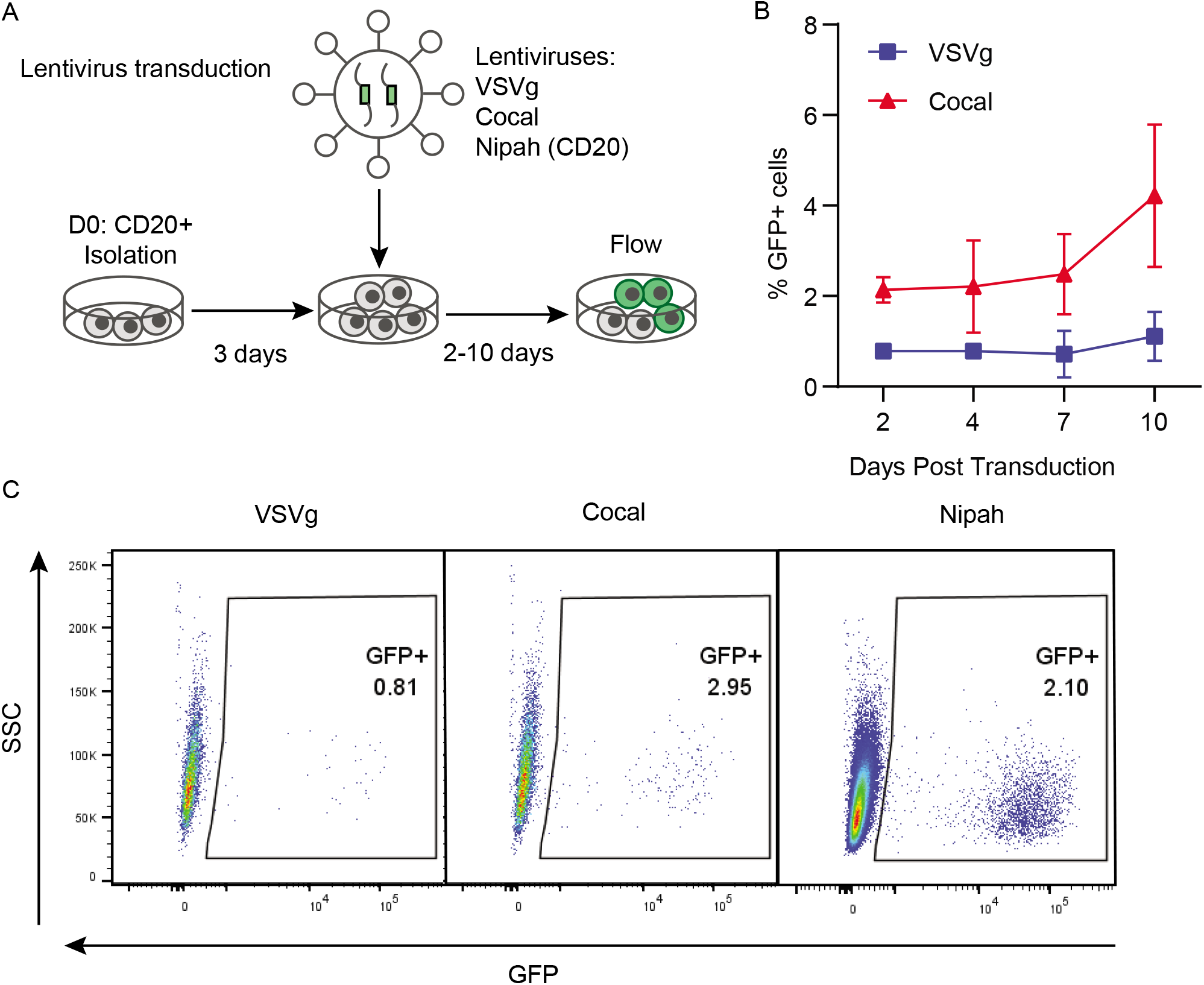
(A) Diagram of the lentiviral transduction and culturing protocol. (B-E) The percent GFP expression was quantified by flow cytometry in CD3^-^CD14^-^ NHP B cells transduced with lentivirus pseudotyped with either VSVg or Cocal envelopes at an multiplicity of infection of 40 (2 donors, n=2) or with a Nipah anti-CD20 pseudotyped virus (1 donor, n=1). (B) Graph showing GFP expression overtime for both VSVg and Cocal Pseudotypes (C-E) Representative flow plots showing GFP percentage in CD3^-^CD14^-^ cells 4 days post transduction with lentivirus pseudotyped with the indicated envelopes.

## DISCUSSION

An important proof of concept experiment for developing a future human plasma cell-based therapy is assessment of plasma cell survival and production capacity in an immunocompetent system. Although NHP models have been used extensively to study the response to vaccines^10^ and plasma cell longevity^11^, we are unaware of reports showing *ex vivo* differentiation and expansion of NHP B cells. To address this challenge, we developed methods to purify, expand, and differentiate *rhesus macaque* B cells *ex vivo*. We achieved a 10-fold expansion of NHP B cells and observed efficient differentiation into plasma cells. In our hands, due to the lack of mAbs that efficiently recognize key primate surface markers, the most efficient surrogate marker for differentiation into antibody-secreting cells is ELISPOT quantification of immunoglobulin secretion. Finally, we identified the serotypes (2.5 and D-J) and conditions necessary for efficient transduction of NHP B cells with AAV vectors for the purposes of producing a secreted exogenous protein (BAFF).

To address the question of how many plasma cells will be required in primates to generate a neutralizing dose of antibody or relevant levels of alternative therapeutic proteins, there are several variables to be considered: half-life of the protein, effective dose, the production of protein per plasma cell and the engraftment capacity of the plasma cell product. Fortunately, for most clinical mAbs, we know the half-life and the required neutralizing dose. For example, the recently developed half-life extended version of palivizumab, MEDI8897, and standard palivizumab have half-lives of approximately 100 days (2400 hours^12^) and 20 days respectively^13^. Given the half-life (t_1/2_) and volume of displacement (V_d_; 880 mL in a 15 kg adult primate subject), one can estimate the rate of drug clearance (CL). Likewise, using the constant rate infusion equation, given a desired steady-state concentration (*C_ss_*), we can calculate the required daily production. The trough concentration required for neutralization of pathogens is known for many mAbs including CR6261 in influenza (^~^10 mg/mL^14^), VRC01 in HIV (^~^10 mg/mL^15^) and MEDI8897 in RSV (^~^8 mg/mL^12^). Applying these equations to MEDI8897, the approximate daily production (*R_0_*) required for RSV neutralization is approximately 56 mg/day. Because fully differentiated long-lived plasma cells are estimated to produce between 50 pg/cell/day^16^, we predict that stable engraftment of 5 million primate plasma cells would be sufficient to produce a prophylactic dose of antibody at steady-state.

Upon successful development of engineered primate plasma cells, we expect that production of a non-clinical cell therapy product could be multiplexed and delivered singly or repeatedly into an autologous donor. recipients. Using leukapheresis, approximately 5 billion nucleated cells can be acquired from a single donor in a single extraction^17^ and ^~^10% of these cells are expected to be B cells. Based on the rates of expansion observed here (10-fold), following expansion and differentiation, we predict that it will be possible to generate ^~^5 billion engraftable plasma cells from a single primate leukapheresis extraction. Given the number of cells we typically transplant into mice to elicit therapeutically relevant and stable antibody secretion^1^, approximately 2.5 billion cells will be required to generate similar antibody levels in an autologous primate transfer. In summary, these results suggest that proof-of-concept experiments using one or more doses of engineered primate plasma cells are likely to be feasible with minor adjustments to the protocol presented here.

## METHODS

### B cell isolation

Frozen rhesus macaque (*macaca mulatta*) PBMCs were obtained from a commercial source (BioIVT). Using an NHP specific CD20 positive selection isolation kit (Miltenyi Biotec; Figure S1), we isolated CD20+ cells.

### B cell culture conditions

Unless otherwise stated, all cells were cultured in a base media composed of Iscove’s modified Dulbecco’s medium (IMDM) (Gibco) supplemented with 10% FBS (Omega Scientific, FB-11), 1% Glutamax (Gibco), and 55 mM beta-mercaptoethanol (BMe). Where noted cells were also cultured in ImmunoCult-XF (ICXF) base media (StemCell, 10981) The commercial expansion cocktail we used was ImmunoCult^™^-ACF Human B Cell Expansion Supplement (StemCell,10974). For the defined B cell expansion and differentiation cocktails, the compositions are as follows. Phase I: 100 ng/mL recombinant human MEGACD40L (Enzo Life Sciences), 1 mg/mL CpG oligodeoxynucleotide 2006 (Invitrogen), 250 ng/mL IL-2 (PeproTech), 50 ng/mL IL-10 (PeproTech), and 10 ng/mL IL-15 (PeproTech), and IL-21 (PeproTech, 200-21). Phase II: IL-2 (250 ng/mL), IL-6 (50 ng/mL, PeproTech), IL-10 (50 ng/mL), and IL-15 (10 ng/mL). Phase III: IL-6 (50 ng/mL), IL-15 (10 ng/mL), and human interferon-a 2B (15 ng/mL, Sigma-Aldrich). All cell counts were done manually using a hemocytometer and trypan blue exclusion.

### Flow cytometry

Flow cytometry was run on an LSR II flow cytometer (BD Biosciences) and data were analyzed using FlowJo software (Tree Star). All cells were fixed using Fix/Perm (BD Biosciences) for 20 min prior to intracellular staining and/or analysis. The flow cytometry antibody panels (Figure S1) and gating schemes (Figures S2-S4) are detailed in supplemental figures.

### ELISpot and ELISAS

Membranes were coated with an anti-human IgG capture antibody (Thermo Fisher Scientific, A18813). Cells were then added to the plate in duplicate in 2 fold serial dilutions and incubated for 24hrs prior to detection using an alkaline phosphatase-linked anti-human IgG detection antibody (Jackson, 109-055-008). Protein secretion levels in culture supernatant were quantified via ELISA for human IgG (Invitrogen, 50-112-8849) and human BAFF (R&D Systems, DY124-05) using protocols recommended by the manufacturer.

### Virus production and transduction

All viruses and all pseudotypes, AAV and lentivirus, were made in house previously published methods. GFP-BAFF was previously published in King et al. AAV transduction was done on day three of culture. First cells were replated at 1.5×10^6^ cells/mL (unless otherwise noted) in IMDM + Bme + Glutamax + commercial supplement without FBS, then virus was added to culture at 20% by volume unless indicated otherwise and incubated at 37°C. After two hours FBS was added back to the media for a final concentration of 10% and returned to 37°C. The next day the media volume was doubled in order to minimize any cell death caused by the addition of the AAV. For Lentivirus transduction, cells were replated at 1.5×10^6^ cells/mL IMDM alone, and virus was added. Cells were spinoculated for 30min at RT at 400xg, then returned to 37°C for six hours. After six hours cells were spun down for 5 min at 400xg, half the media volume was taken, and IMDM was added back with two times the concentration of FBS, BMe, Glutamax, and commercial supplement and returned to 37°C.

### Single Cell RNA Sequencing

Single cell cDNA libraries were created using the 10x genomics platform (3’ chemistry V3.1 dual index kit, 10x genomics). And preliminary sequencing was done on an Illumina MiSeq machine using a MiSeq Reagent Kit v3 300-cycles. Data was processed and interpreted using Cell Ranger and Loupe Browser (10x genomics)

