## Supplemental Figures for "Efficient methods for generation and expansion of, and gene delivery to Rhesus Macaque plasma B cells"

Figure S1

| Reagent | Company | Cat |
|---|---|---|
| NHP CD20 Isolation Kit | Miltenyi | 130-091-105 |
| IMDM | Gibco | 12-440-053 |
| ICXF | StemCell | 10981 |
| StemCell B-cell supplement ACF | StemCell | 10974 |
| MEGACD40L | Enzo Life Sciences | |
| CpG oligodeoxynucleotide 2006 | Invitrogen | |
| IL-2 | PeproTech | AF-200-02 |
| IL-10 | PeproTech | |
| IL-15 | PeproTech | |
| IL-21 | PeproTech | 200-21 |
| IL-6 | PeproTech | |
| Human interferon-a 2B | Sigma-Aldrich | |
| APC-Cy7 anti-CD3 clone:SP34-2 | BD Bioscience | 557757 |
| A700 anti-CD3 clone:SP34-2 | BD Bioscience | 557917 |
| BV605 anti-CD4 clone:L200 | BD Bioscience | 562843 |
| APC.Cy7 anti-CD14 clone:MoP9 | BD Bioscience | 557831 |
| PerCPCy5.5 anti-CD20 clone:L27 | BD Bioscience | 340955 |
| BV605 anti-CD31 clone:WM59 | Biolegend | 303122 |
| PE anti-CD31 clone:WM59 | BD Bioscience | 555446 |
| mFluor450 anti-CD38 clone:OKT10 | Caprico Biotechnologies | 1008144 |
| PE anti-CD56 clone:MY31 | eBioscience | 12-0569-71* |
| PE anti-CD138 clone:clone | eBioscience | 63-1389-41* |
| AF700 anti-IgG clone:G18-145 | BD Bioscience | 561296 |
| FITC anti-IgM clone:G20-127 | BD Bioscience | 555782 |
| IgG capture antibody (ELISpot) | Invitrogen | A18813 |
| IgG AP detection antibody (ELISpot) | Jackson | 109-055-008 |
| Human IgG ELISA | | |
| Human BAFF DuoSet ELISA | R&D Systems | DY124-05 |

**Panel 1: Day 0 Purity assessment**

| Antigen | Fluor | Clone | Cat# | Dilution |
|---|---|---|---|---|
| Live Dead | UV/ A350 | N/A | | 1:1000 |
| CD20 | PerCPCy5.5 | L27 | 340955 | 1:20 |
| CD14 | APC.Cy7 | MoP9 | 557831 | 1:200 |
| CD56 | PE | MY31 | 12-0569-71 | 1:200 |
| CD4 | BV605 | L200 | 562843 | 1:200 |
| CD3 | A700 | SP34 | 557917 | 1:200 |

**Panel 2: Phenotyping and transduction efficiency panel**

| Antigen | Fluor | Clone | Cat# | Dilution |
|---|---|---|---|---|
| Live Dead | UV/ A350 | N/A | | 1:1000 |
| CD20 | PerCP-Cy5.5 | L27 | 340955 | 1:100 |
| CD14 | APC.Cy7 | MoP9 | 557831 | 1:200 |
| CD31 | PE | WM59 | 555446 | 1:100 |
| CD38 | mFluor450 | OKT10 | 1008144 | 1:100 |
| CD3 | AF700 | SP34 | 557917 | 1:200 |
| Transduction | GFP | -- | --- | --- |

**Panel 3: Phenotyping and intracellular Ig staining**

| Antigen | Fluor | Clone | Cat# | Dilution |
|---|---|---|---|---|
| Live Dead | UV/ A350 | N/A | | 1:1000 |
| CD20 | PerCP-Cy5.5 | L27 | 340955 | 1:20 |
| CD14 | APC-Cy7 | MoP9 | 557831 | 1:200 |
| CD138 | PE | DL-101 | | 1:100 |
| CD38 | mFluor450 | OKT10 | 1008144 | 1:50 |
| CD3 | APC-Cy7 | SP34 | 557917 | 1:200 |
| CD31 | BV605 | WM59 | 303122 | 1:100 |
| IgG-ic | AF700 | G18-145 | 561296 | 1:200 |
| IgM-ic | FITC | G20-127 | | 1:200 |

A

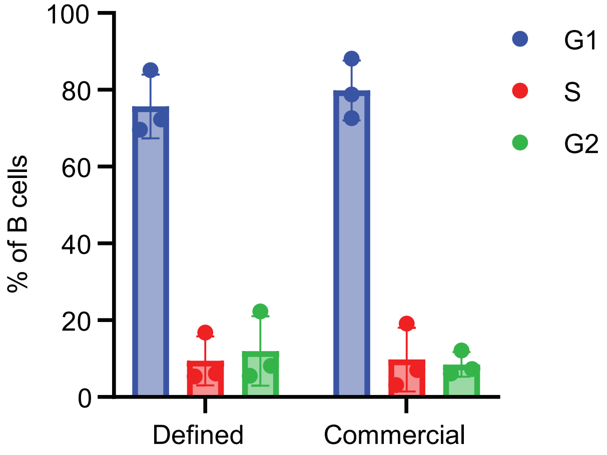


Supplementary Figure 2: (A) NHP B cells were cultured for 3 days in either the defined or comercial medias, and on day 3 flow with propidium iodide was run to quantify cells in different cell cycle phases.

**A**

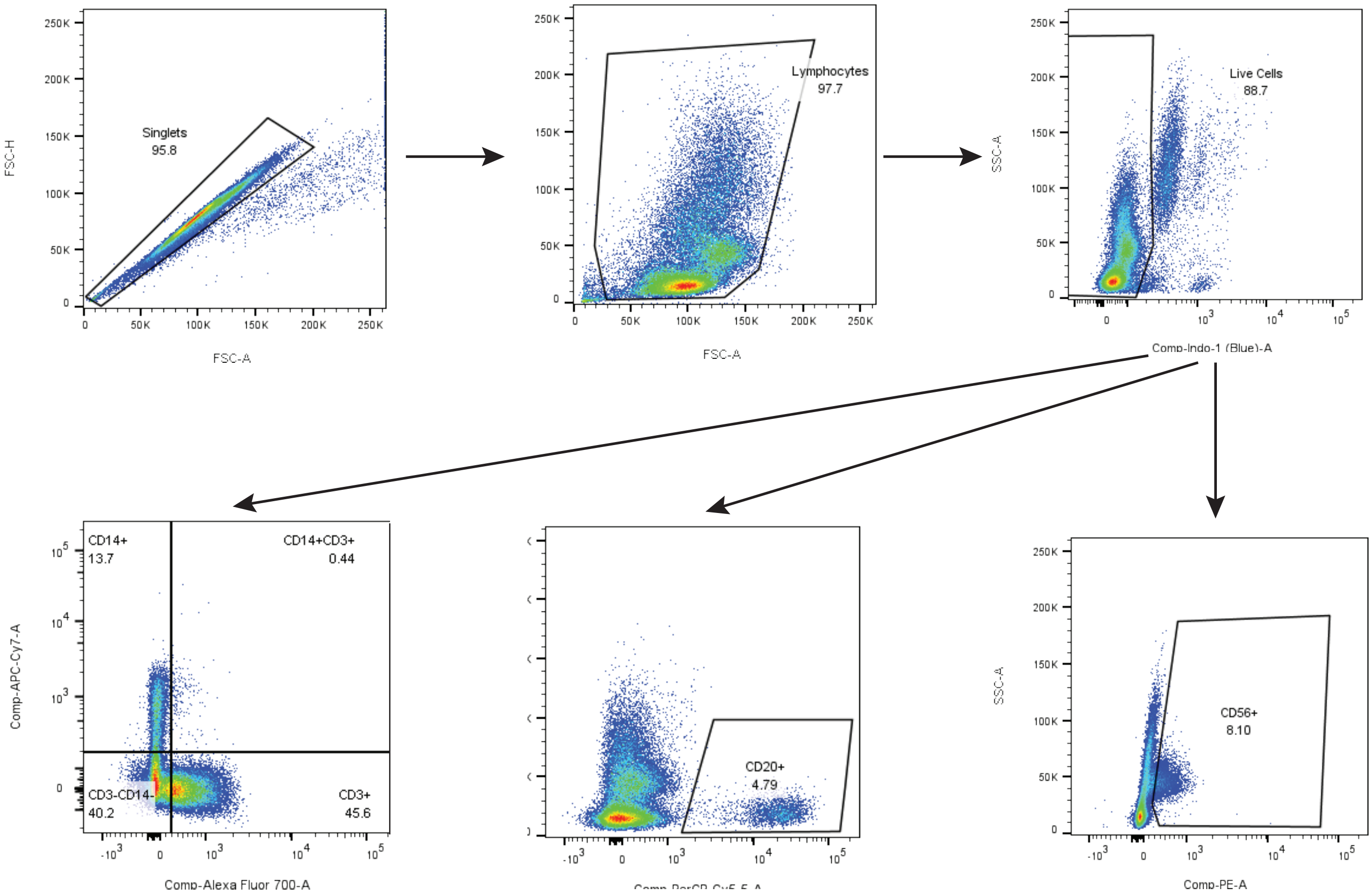


Supplementary Figure 3: (A) Gating scheme for flow cytometry Panel 1 (S1B) to determine purity of at the time of isolation

A

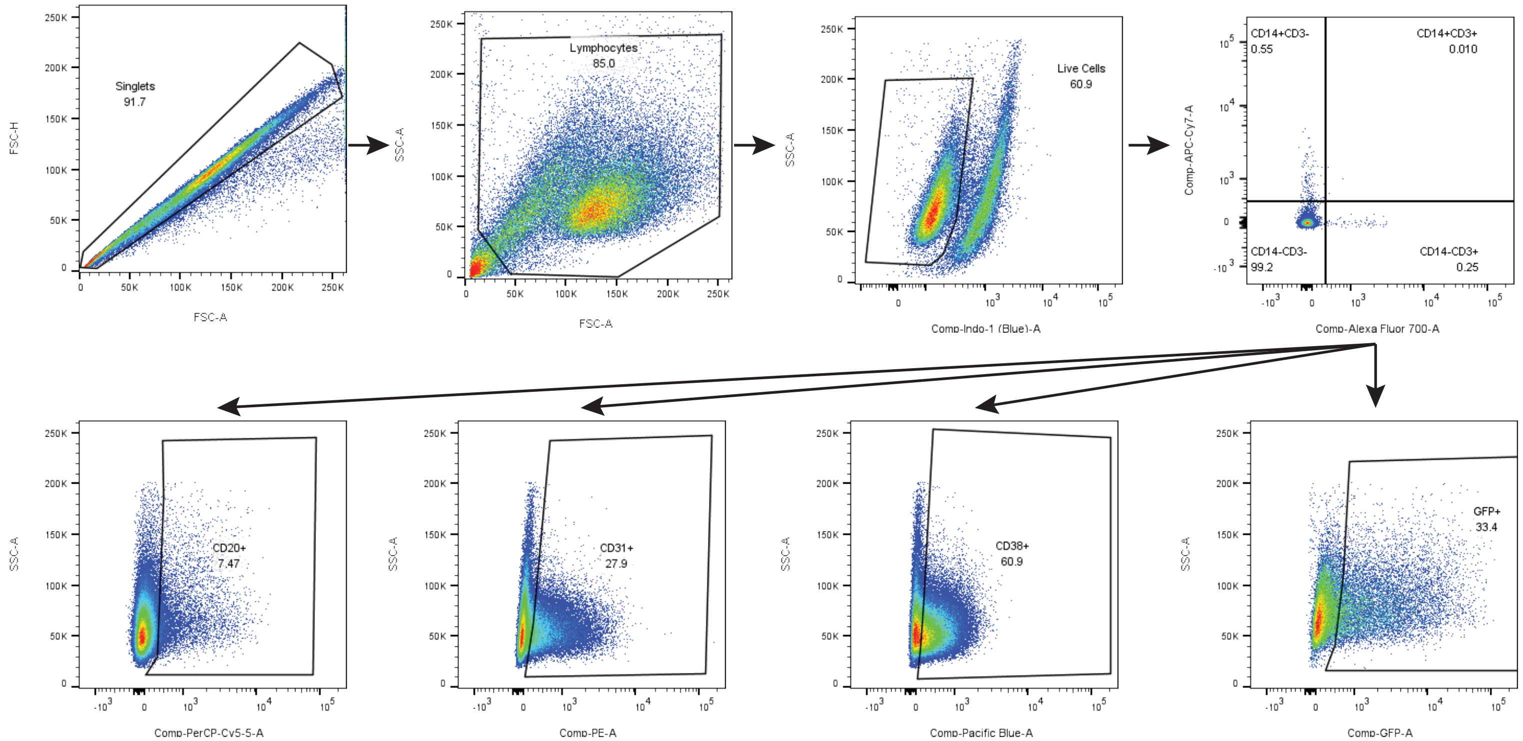


Supplementary Figure 4: (A) Gating scheme for flow cytometry Panel 2 (S1) to determine B cell phenotype and transduction efficiency through GFP expression

A

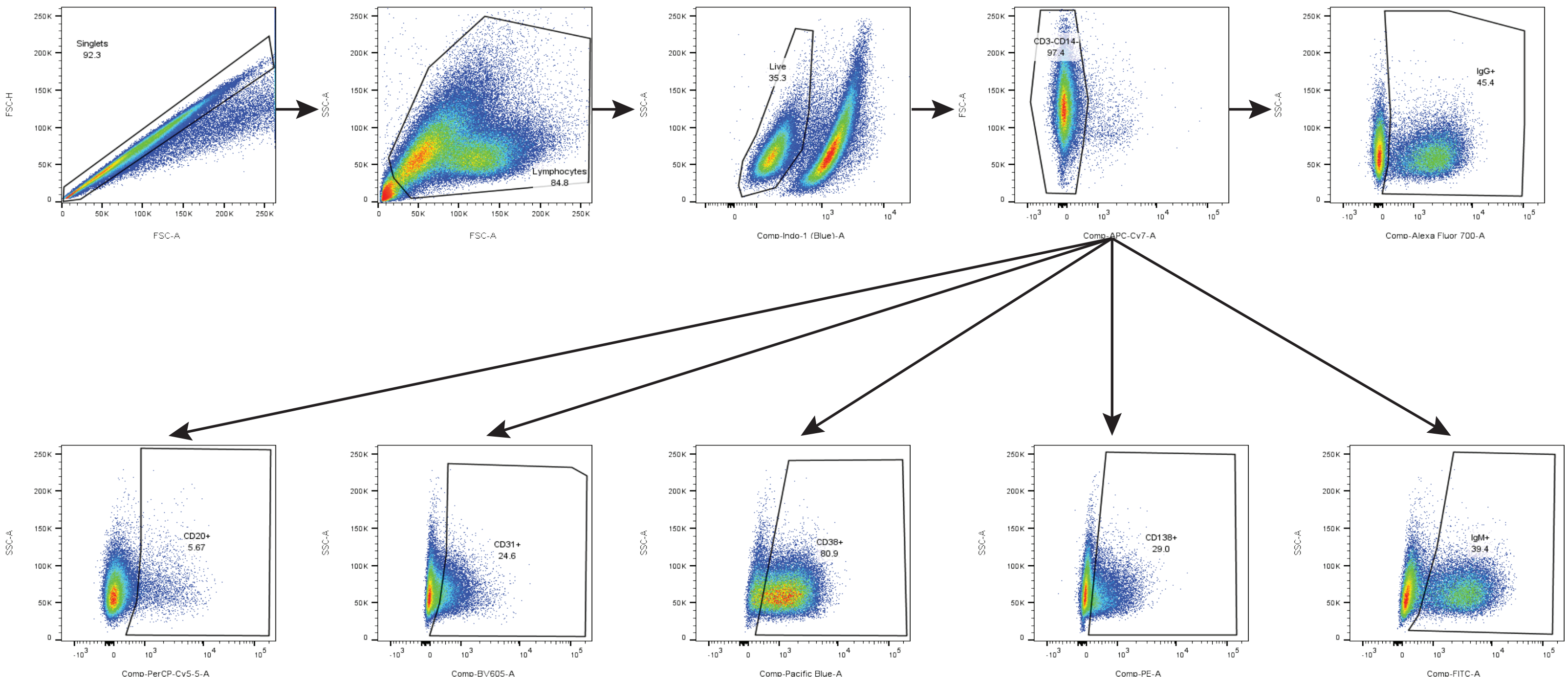


Supplementary Figure 5: (A) Gating scheme for flow cytometry Panel 3 (S1) to determine B cell phenotype and intracellular Ig production

A

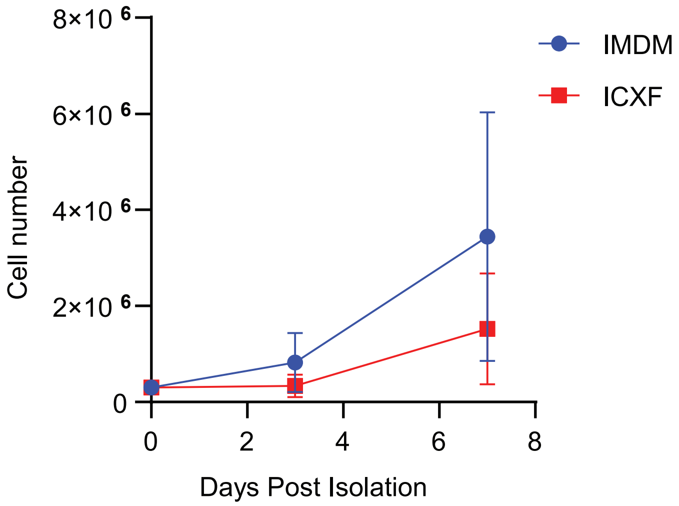


Supplementary Figure 6: (A) NHP B cells were cultured in two different basse mediums IMDM and ICXF at a low concentration (plated at 2.5x10cells/mL, and maintained at 1x10$^{5}$ cells/mL). Cells were counted on days 3 and 7.

A

B

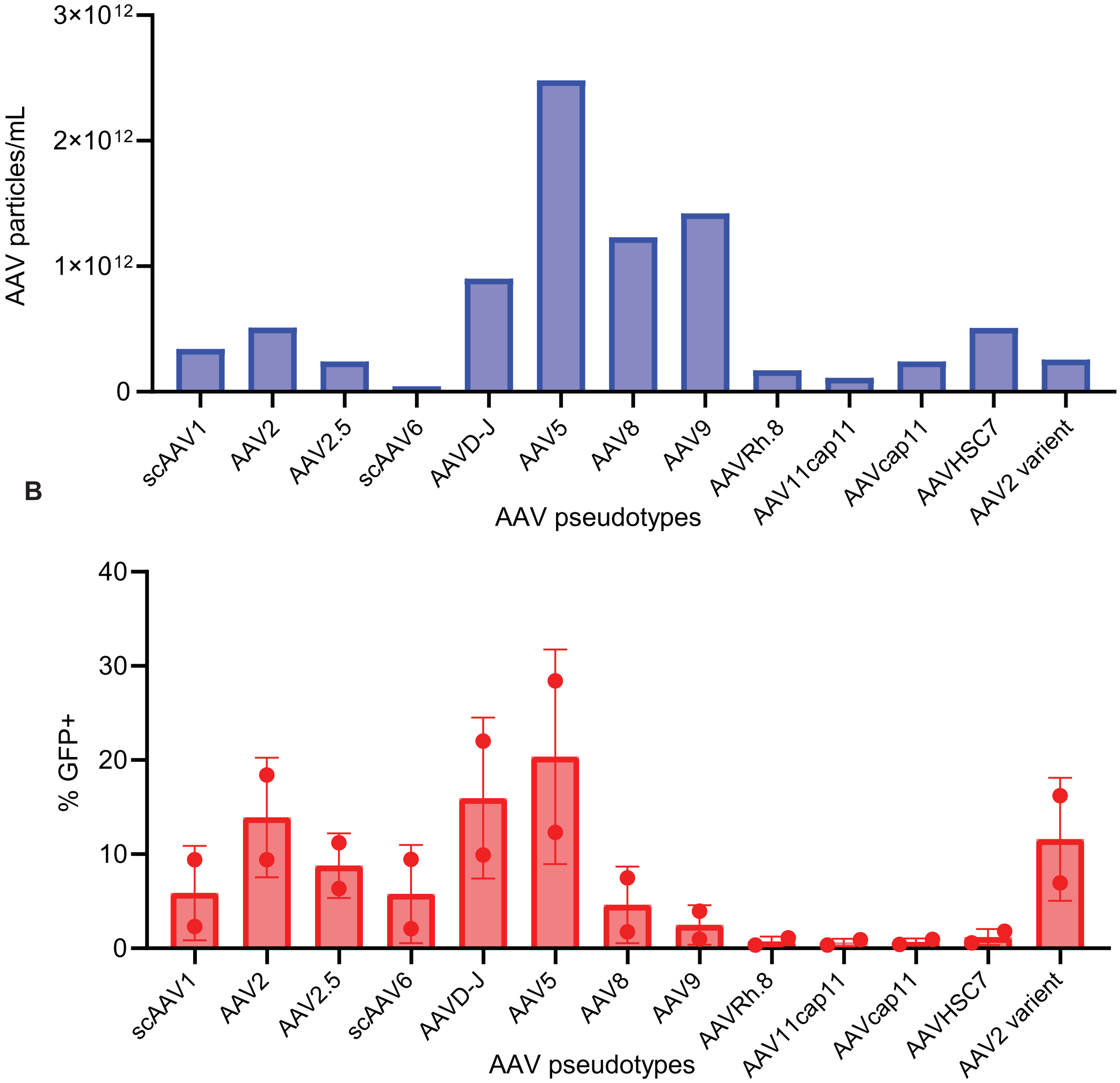


Supplementary Figure 7: (A-B) NHP B cells were transduced on day 3 of culture. 20% by volume was added of GFP cancapsulated in different AAV pseudotypes, with varrying titers (A), and flow for GFP (B) was run 2 days later.
